# The Effect of Perturbation Variability on Sensorimotor Adaptation Does Not Require an Implicit Memory of Errors

**DOI:** 10.1101/2022.05.30.493844

**Authors:** Tianhe Wang, Guy Avraham, Jonathan S. Tsay, Richard B. Ivry

**Author notes:** Corresponding author: Tianhe Wang.

## Abstract

In a recent paper^1^ entitled, “An implicit memory of errors limits human sensorimotor adaptation” Albert and colleagues presented a model in which the adaptive response of the sensorimotor system is flexibly modulated by recent experience, or what they refer to as a “memory of errors”. This hypothesis stands in contrast to prevailing models in which automatic and implicit responses to movement errors are relatively insensitive to the statistical properties of the environment^2–6^. A prime example of this rigidity is that the adaptation system exhibits a saturated response to large errors, resulting in a non-linear motor correction function, a feature that is independent of experience^4,5,7^. Here we show that the key results reported in Albert et al. are fully explained by presupposing this rigid “motor correction” function without reference to memory-dependent changes in error sensitivity. As such, the evidence presented in Albert et. al. does not support the claim that the history of errors modulates implicit adaptation.

## Results

A memory of experienced errors would be computationally advantageous, allowing the system to calibrate an optimal response. For example, the responsiveness of the system could be enhanced in stable environments where similar errors are experienced across trials. To test this hypothesis, Albert et al. rotated the visual feedback by 30° relative to true hand position as human participants reached to a target, comparing conditions in which the mean rotation had high variance (12° S.D.) or zero variance (0° S.D.). Across a series of experiments, implicit adaptation was attenuated in the high variance condition. To account for this finding, the authors presented a state-space model in which the learning rate is modulated in a memory-dependent manner. In their implementation, sensitivity increases with repeated exposure to errors with the same sign and decreases when the sign reverses.

In developing their model, Albert et al. assumed that the function describing error sensitivity is initially flat, changing its shape as a function of short-term experience. However, an extensive literature using a wide range of methods has shown that error sensitivity decreases with error size (Fig. 1b)^4,6,8^. Correspondingly, the motor correction function, describing the error-dependent trial-to-trial change in the sensorimotor map, increases over a range of small errors up to the saturation point and eventually decreases for very large errors (Fig. 1a). From a Bayesian inference perspective, this function reflects the discounting of large errors^9^; alternatively, this function may reflect the upper limits of plasticity in the sensory^10^ or motor system^5^. Importantly, in either scenario, the motor correction function is assumed to be a “primitive” or an established prior^9^, existing independent of the history of errors. In this commentary, we show that a fixed non-linear motor correction function is sufficient to account for the results presented by Albert et al. Importantly, this alternative account does not include short-term, experience-dependent changes in error sensitivity.

**Figure 1.**
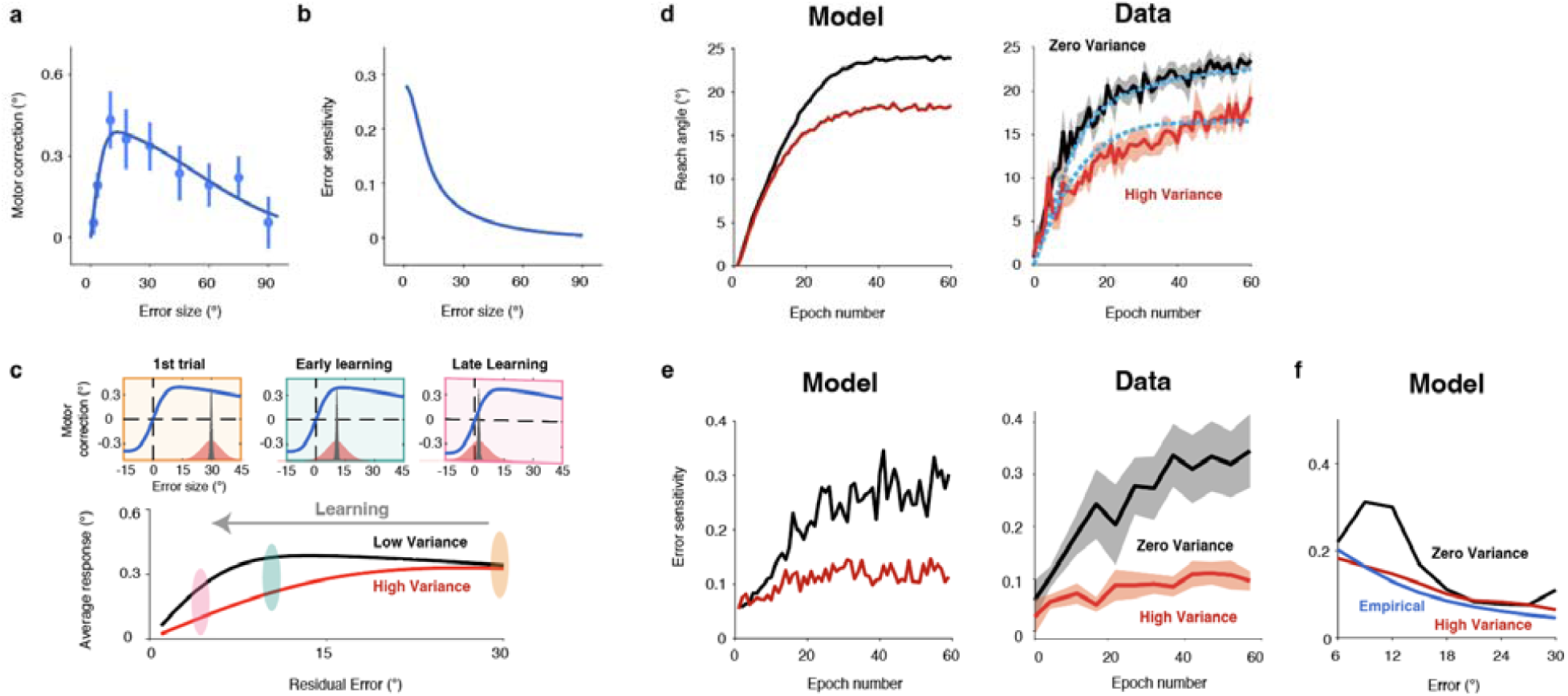
A non-linear motor correction function is sufficient to explain the effect of perturbation variability. **a**, Changes in hand angle from trial n to n + 1 as a function of the error size on trial n (Tsay et al., 2021). We used these data to estimate the motor correction function given its relatively high sampling density. Dots represent group median and bars represent the standard error of the means. Solid line denotes the best-fitting model. **b**, Error sensitivity, defined as the motor correction divided by error size, reduces monotonically as error size increases. **c**, Perturbation variability (red, high variability; black, low variability) impacts the distribution of the experienced errors and, thus influences the distribution of motor corrections (upper panel). When the error is large at the start of learning, the average response is similar between low and high variance (lower panel). However, as learning unfolds, the error distribution for the high variance condition will span the concave region of the motor correction function, resulting in attenuation of the average motor correction. **d-e**, Time course of implicit adaptation **(d)** and error sensitivity **(e)** in Experiment 6 of Albert et al. The results from the experiment are graphed on the left of each panel and the results from our simulations using the fixed motor correction function shown in **(a)** are presented on the right. Shaded areas represent standard error. The blue dashed lines in **(d)** depict the simulated functions derived from the memory of errors model proposed by Albert et al. Note that we focused on Exp 6 since explicit (aiming) processes contribute to performance in the other experiments in their paper. **f**, Recovered error sensitivity functions from simulations using an estimate of motor noise from Albert et al. The functions fail to recover the error sensitivity function used in the simulations (blue curve from 1b, same for both conditions).

For our simulations, we used a motor correction function derived from a task in which the feedback varied from trial to trial (Fig. 1a)^8^. We first examined the consequences of perturbation variability across trials under the assumption that the motor correction function is fixed (Fig. 1c). Perturbation variability will impact the probability distribution of the experienced errors, and due to the non-linearity of the motor correction function, can impact the average motor correction. If the experienced errors all fall within the linear zone (Fig. 1c, 30° error), the average motor correction will be the similar for perturbation conditions involving either low or high variability. However, if the experienced errors span the concave region of this function (Fig. 1c, 10° error), high variability will result in an attenuated average response. This attenuation occurs since averaging across the corrections to errors smaller than the mean error are no longer balanced by corrections to errors larger than the mean error.

To verify that a fixed motor correction function can capture the variability effects reported in Albert et al., we simulated the learning functions for a visuomotor adaptation task using perturbations centered at 30° with either high variability (12° S.D.) or zero variability (0° S.D.; Experiment 6 in Albert et al). We used a classic state-space model with the trial-to-trial update determined by an empirically-derived motor correction function (Fig 1a)^8^ and the retention factor reported in Albert et al. (see Supplementary Methods). The simulated results capture the key features of their behavioral results: The high variance condition results in a slower learning rate and larger residual error (i.e., lower asymptote, see Fig. 1d). Moreover, the simulations capture the time-course or the error sensitivity functions (Fig 1e). There is a small difference in error sensitivity early in learning when the experienced errors fall in the descending linear zone of the error correction function. However, the functions diverge as learning progresses when the experienced errors encompass the non-monotonic region. This simulation results do not depend on the specific shape of the motor correction function: The same pattern of results can be obtained using a wide range of functions in which the motor correction function either decreases or saturates for large errors (Extended Data Figs. 1-2).

We note the memory of errors model of Albert et al. suggests a mechanism in which the error sensitivity function emerges with experience. Namely, error sensitivity will be lower for large errors simply because these are experienced less frequently than small errors. Correspondingly, their model predicts a motor correction function that bears a non-linear shape within a limited range (0-30°; Extended Data Fig. 3a), independent of how the function is initialized. However, rather than view the motor correction function as the end result of a learning process, multiple lines of evidence suggest that this function exists independent of the errors that participants experience in an experimental session. First, error sensitivity varies with error size in the initial responses to a perturbation^4,5^. Second, the non-constant error sensitivity function (i.e., non-linear motor correction function) is seen in tasks that randomly vary the sign and size of the perturbation from trial to trial^7–9,11^. In contrast, the Albert et al. model predicts a constant error sensitivity function (a linear motor correction function, Extended Data Fig. 3c) under a random perturbation schedule since the enhancing effect from the repeated exposure to errors of a given sign will be nullified by the attenuating effect from sign reversals that occur with equal frequency. Third, the motor correction function is not affected by the probability of the sign reversals^7,12^, a result at odds with the prediction of the Albert et al. model.

We note that one analysis reported by Albert et al. appears to be inconsistent with our memory-free model: The error sensitivity function is higher for the zero variance condition even when the data are binned to “equate” error size between the two conditions (Fig. 4a in the original paper). However, given the presence of motor noise, the estimation of the error sensitivity function cannot be recovered when the perturbation is fixed (i.e., blocked design). We failed to recover the error sensitivity functions in the simulations when using an estimate of motor noise (3° SD) based on the baseline phase in Albert et al. (Fig. 1f; Extended Data Figs. 4). Moreover, the recovered error sensitivity function is higher in the zero-variance condition despite the fact that the same underlying function was used in both conditions in the model.

We recognize that our simulation results do not refute the possibility that the response of the implicit motor adaptation system may be modulated by recent experience. That is, a non-linear sensitivity function established from long-term experience might undergo transient modification in response to recently experienced errors. However, the simulations presented in this commentary demonstrate that a short-term memory process is not needed to explain the behavioral differences that emerge as a function of perturbation variability. Future work is needed to disentangle the contribution of an a priori, non-linear motor correction function and effects associated with contextual factors such as recent error history.

## Data availability

The code that supports this commentary is available at https://osf.io/exg4c/

## Author contributions

All four authors contributed to the conceptual development of this project. T.W. performed the model simulations and wrote the initial draft of the paper, with all of the authors involved in the editing process.

---

**Extended Data Fig. 1.**
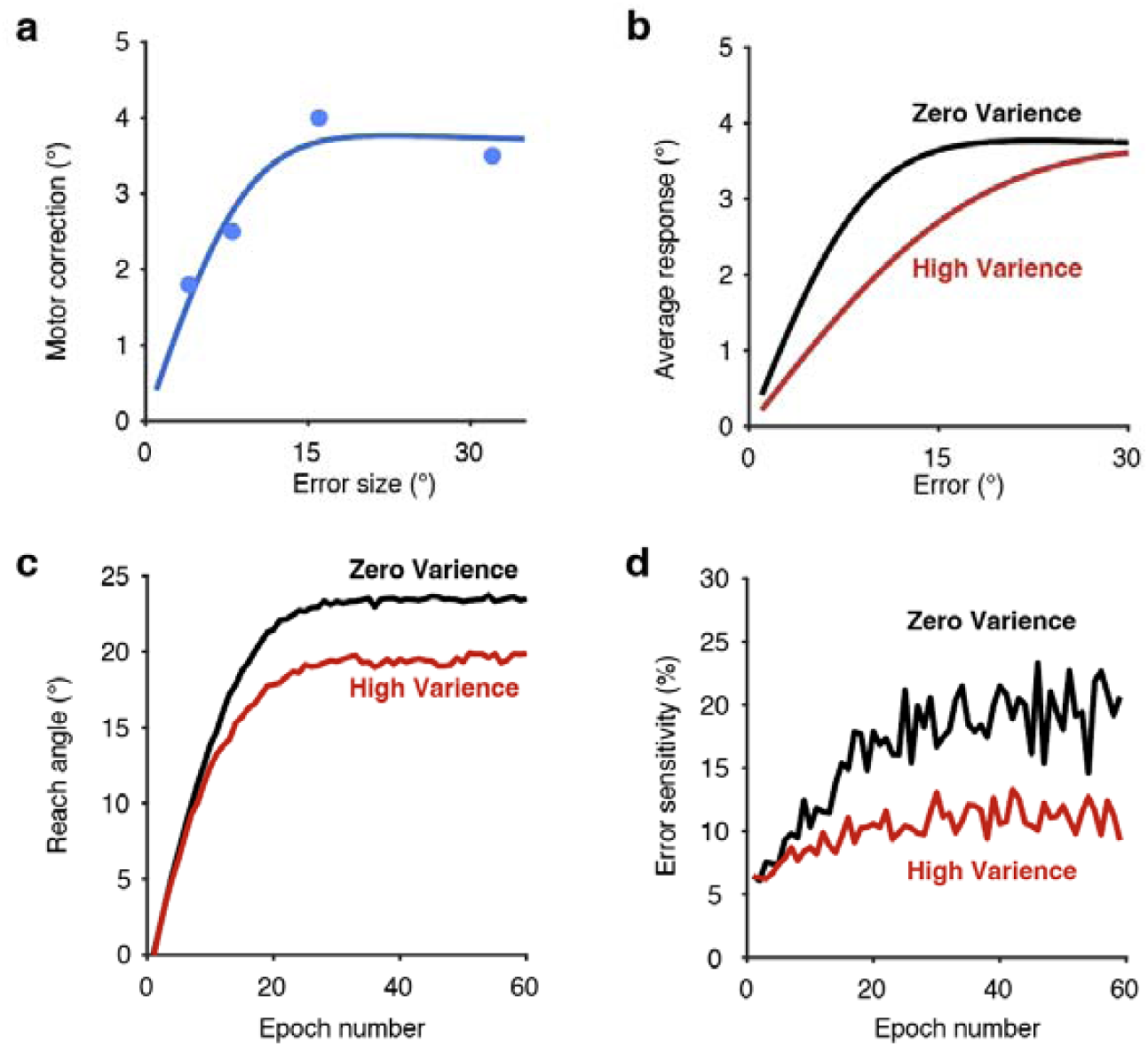
Replication of Figure 1 using a motor correction function derived from Hutter and Taylor, 2018. **a**, Motor correction as a function of the error size. Dots correspond to the mean values in Exp. 2 of Hutter & Taylor. The solid line denotes the best-fit model. **b**, Average response in the low (black) and high (red) perturbation variability conditions. **c-d**, Time course of implicit adaptation **(c)** and error sensitivity **(d)** from simulations using our model.

**Extended Data Fig. 2.**
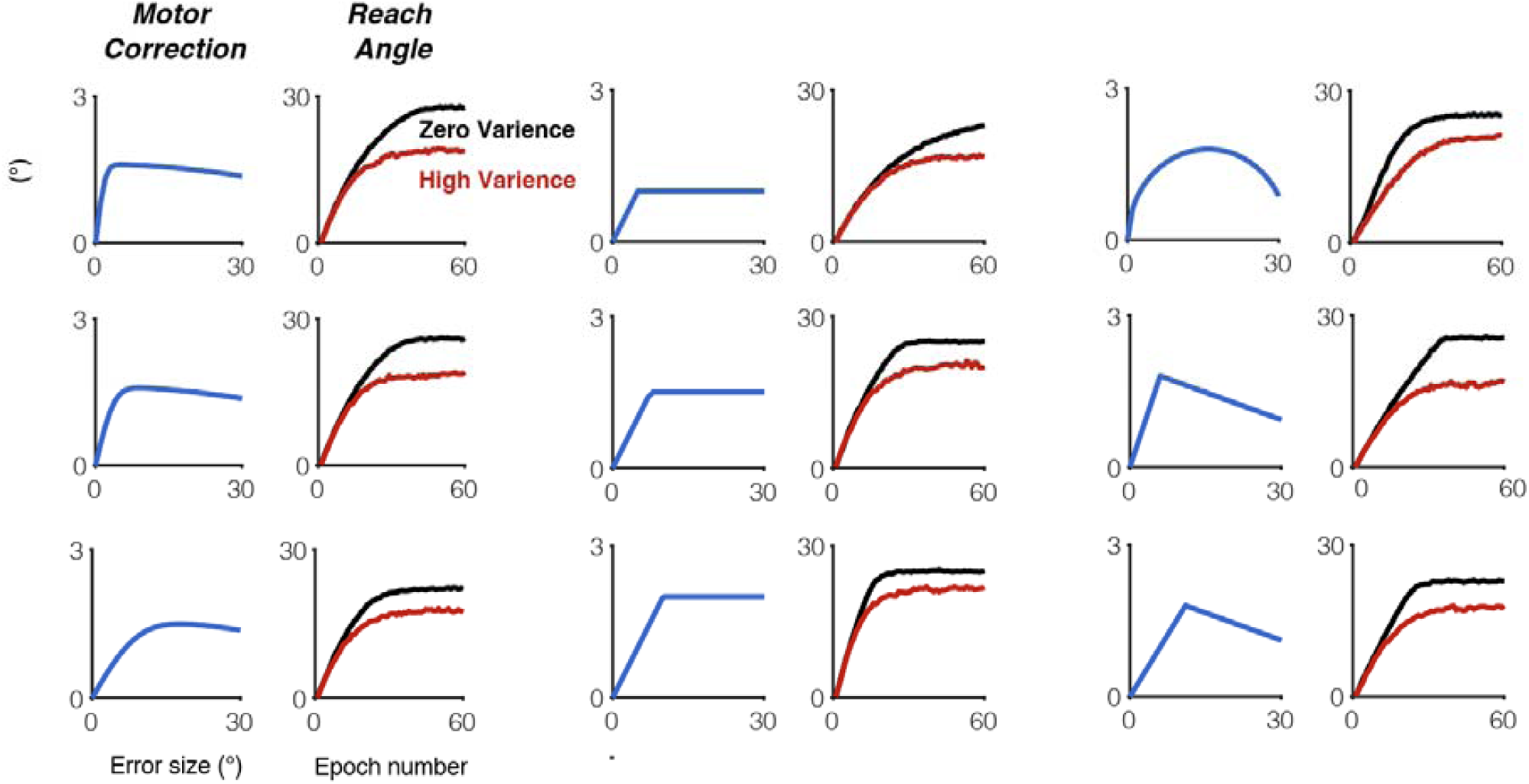
Effect of perturbation variability persists with a variety of hypothetical motor correction functions. For each subplot, the motor correction function is depicted on the left and the time course of the learning function is depicted on the right. The concave motor correction function either saturates or decreases for large errors. We varied the curvature and the peak position of the functions. All of the tested motor correction functions result in attenuated implicit adaptation for the high variability condition (red) relative to the zero-varience condition (black).

**Extended Data Fig. 3.**
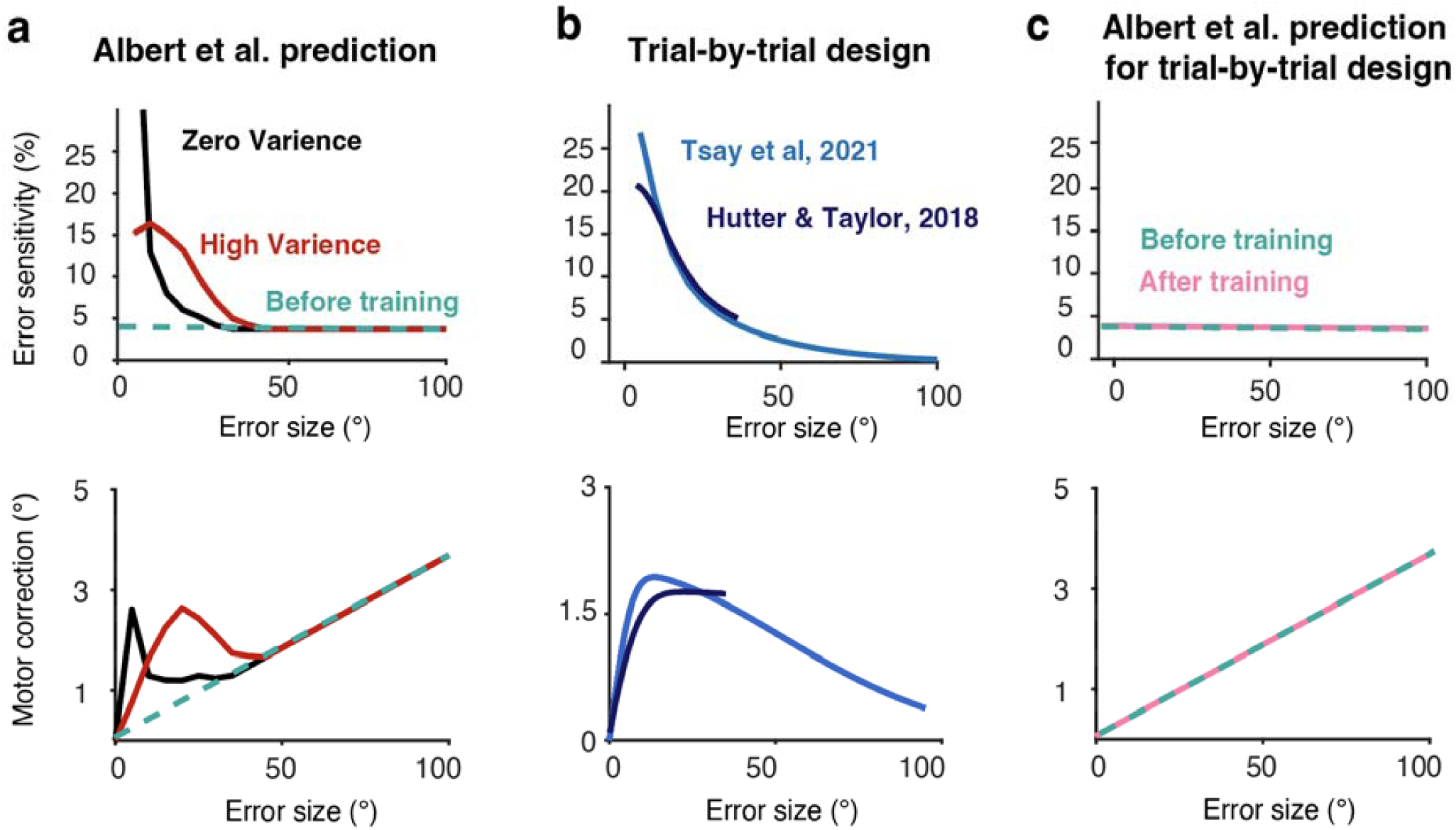
The memory of errors model fails to predict the error sensitivity function observed in designs with randomized perturbations. **a)** Albert et al. assumed that the error sensitivity function is initially a constant (top), resulting in a linear motor correction function (bottom). By itself, this function fails to capture the effect of perturbation variability. In their model, experiencing consistent or inconsistent errors will respectively, increase or decrease error sensitivity. This memory process will result in a non-constant error sensitivity function (top) and a non-linear motor correction function (bottom) for error sizes that were experienced during the experiment (0 to 30°). Note that for larger errors, the model predicts constant error sensitivity and thus a linear motor correction function (although in their experiments, errors of this size would be very infrequent). b) The error sensitivity function (top) and the non-linear motor correction function (bottom) estimated from Tsay et al (2021) and Hutter and Taylor (2018) in which the sign and size of the perturbation was randomized across trials. c) The memory of errors model fails to replicate the error sensitivity functions as measured by randomized perturbations. Instead, it predicts that error sensitivity (top) will remain constant and that the motor correction function will remain linear (bottom) after training.

**Extended Data Fig. 4.**
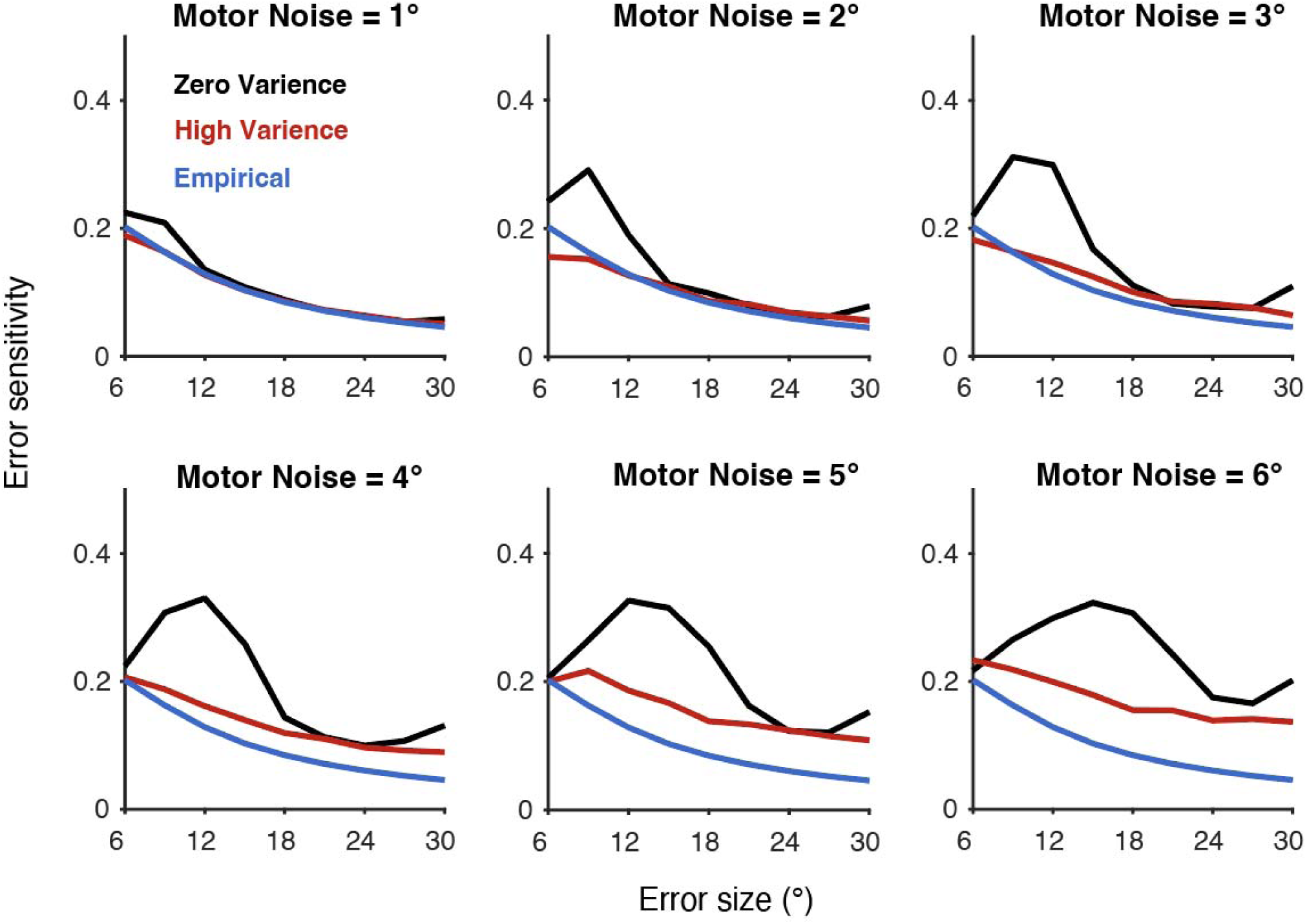
The error sensitivity function is not recoverable in the fixed perturbation design used by Albert et al. Recovered error sensitivity functions from simulations using motor noise sampled from a normal distribution with different standard deviations (range: 1-6°). It becomes more and more difficult to recover the error sensitivity function as the motor noise increases. The recovered error sensitivity function is always higher in the zero-variance condition.

## Supplementary information

### Supplementary Methods

The focus of this note is to ask if the effect of perturbation variability on implicit sensorimotor adaptation can result from a non-linear motor correction function that exists as a prior and is experience-independent. The following provides details of the simulations used to test this hypothesis.

#### Assessing the motor correction function

For the motor correction function (i.e., the prior), we draw on data obtained in previous studies that measured the trial-by-trial response of the adaptation system to a range of error sizes. The primary data set was taken from Tsay et al. (2021) since it had a large range and relatively high sampling resolution. Non-contingent visual feedback was used in that study, a method designed to isolate implicit visuomotor adaptation^3^. The participant reached to a visual target and a feedback cursor was presented at the radial distance of the target but with an angular displacement defined relative to the target rather than the participant’s hand position. This displacement varied randomly across trials (19 conditions: 0°, ± 1.5°, 3.5°, 10°, 18°, 30°, 45°, 60°, 75°, 90°). Participants were fully informed that they had no control over the location of the feedback and should ignore it, always attempting to reach directly to the target. With these instructions, trial-by-trial changes in reaching behavior are implicit^12^. To establish the generality of our findings, we repeated the simulations with a motor correction derived from the data in Hutter and Taylor (2018). In that study, the implicit component of learning was estimated by regressing out the contribution of strategy use as obtained from direct reports (Extended Data Fig. 1). We also performed simulations with a variety of hypothetical motor correction functions (Extended Data Fig. 2).

To obtain a continuous motor correction function, we fit the empirical data with a model that encompasses features of two models for implicit learning: Visual uncertainty^8^ and relevance inference^9^. The visual uncertainty model captures the increase in the size of the motor correction as error size increases for a range of small errors (Tsay et al. 2021). The model assumes that the average motor correction for a given error size depends on the probability, *p*(*positive* | *e*), that the participant perceived the error in the correct direction:

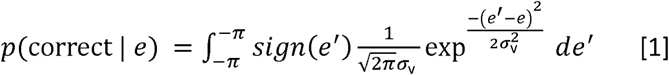

where *sign*(*e*′) indicates the sign of perceived error, *e*′, and *σ* ^*v*^ is the noise in the visual system.

The relevance estimation model captures the decrease in the size of the motor correction for large errors based on the assumption that large errors are likely to be attributed to sources independent of the motor system. Following the derivation of Wei and Kording (2009), error relevancy, *p*(*relevant* | *e*), is determined by the size of the error signal:

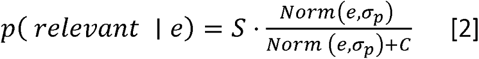

where *Norm*(*e, σ*_*p*_) is the probability of observing e from a normal distribution centered at 0 with a standard deviation of *σ*_*p*_. S and C are scaling factors which, jointly represent the prior belief of how likely an error is related to the motor system.

Taken together, the above features were used to derive a motor correction function:

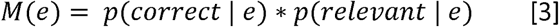

where e is the error size and *M*(*e*) is the motor correction function. We obtained the motor correction function by fitting the combined model with the median error corrections^7,8^, minimizing the residual square error. We opted to use this relatively complex model because simple models (polynomial function, visual uncertainty model alone, relevance inference model alone) are unable to capture both the rapid rise and gradual decline of the motor correction function.

To calculate the error sensitivity function, *z*(*e*), the motor correction function was normalized by *e*:

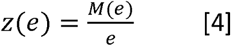

To construct the average motor correction function for the high variance condition, *M*_*high*_ (*e*), and low (zero) variance condition, *M*_*low*_ (*e*), we computed the product of the motor correction function, *M*(*e*), and the perturbation distribution, *norm*(*e*- *e* ′, *S. D*.), and then integrated over *e*′ :

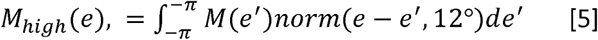

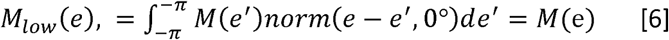

Following Albert et al., we set the standard deviations to 12° and 0° in the high and low variance conditions, respectively.

#### State-space model

To model learning, we employed a standard version of a state-space model:

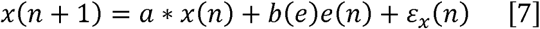

where *x* is the internal estimate of the motor state (i.e., hand movement required to compensate for the perturbation), *a* is the retention factor, *e*(*n*) is the size of the perturbation in trial *n*, and *b* is error sensitivity of a given error size, *e*. *ε*_*x*_ represents planning noise.

The actual motor response on trial *n* is given as:

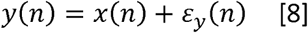

where *y* is the reaching direction relative to the target, determined by *x*(*n*) and the motor noise, *ε*_*y*_.

Note that whereas error sensitivity, *b*, is modulated by experience in the Albert et al. model, it is fixed across trials in our model, only determined by *e*:

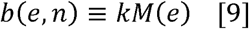

where *k* is a constant that scales the error sensitivity function (see details below).

#### Model simulations

For model simulations, we either used a motor correction function, *M*(*e*), fitted from Tsay et al (2021, Fig. 1) or Hutter and Taylor (2018, Extended Data Fig. 1). While the shapes of those functions are similar, the overall magnitude of the motor correction differs between the studies. This is not surprising since the experimental apparatus and paradigms used to measure implicit adaptation were very different across studies. For example, in Tsay et al., feedback was presented only at the target distance and after the target disappeared, two factors that lead to an attenuated response^14^. For this reason, we scaled the error sensitivity function such that the sensitivity for a 30° error (the initial error experienced at perturbation onset in Albert et al.) was the same across studies. This was done by setting *k*= 5 for the function derived from Tsay et al. and *k*= 0.4 for the function derived from Hutter and Taylor. The retention factor *a* was set as 0.945, a value obtained from the washout phase of Experiment 6 in Albert et al. We assume that learning is the same for reaches to each of the four targets and, thus only simulated the learning of one target for each participant. We dropped the ε_*x*_. The motor noise, *ε*^*y*^ were sampled from normal distributions with a mean of 0 and a S.D. of 3. We note that for motor variability, we only include motor noise measured from the baseline section. This might underestimate the motor variability since the motor variance for individual participant during the training is much larger compared with the prediction of model, perhaps because of active exploration. Consistent with the experimental design of Albert et al., the perturbation in the zero-variance condition was always set to 30°. The perturbations in the high variance condition were randomly sampled from a normal distribution with a mean of 30° and S.D. of 12°. The displayed simulation results were the average of the simulated behavior of 100 participants for each perturbation variance condition.

